# From the Modern Synthesis to the Molecular Synthesis: updating how we teach and assess evolution by natural selection

**DOI:** 10.1101/2021.07.19.452979

**Authors:** Matt Sievers, Connor Reemts, Katherine J. Dickinson, Joya Mukerji, Ismael Barreras Beltran, Elli J. Theobald, Vicente Velasco, Scott Freeman

## Abstract

Evolution by natural selection is recognized as both the most important concept in undergraduate biology and the most difficult to teach. Unfortunately, teaching and assessment of evolution have been impaired by legacy approaches that focus on Darwin’s original insights and the Modern Synthesis’ integration of Mendelian genetics, but ignore or downplay advances from what we term the Molecular Synthesis. To create better alignment between instructional approaches and contemporary research in the biosciences, we propose that the primary learning goal in teaching evolution should be for students to connect genotypes, phenotypes, and fitness. To support this approach, we developed and tested assessment questions and scoring rubrics called the Extended Assessing Conceptual Reasoning of Natural Selection (E-ACORNS) instrument. Initial E-ACORNS data suggest that after traditional instruction, few students recognize the molecular synthesis—prompting us to propose that introductory course sequences be re-organized with the molecular synthesis as their central theme.

## Introduction

When the Modern Synthesis incorporated insights from Mendelian genetics into Darwin and Wallace’s theory of evolution by natural selection, it resolved decades of confusion about the nature of heritable variation as traits change over time. One of the Modern Synthesis’ architects, Theodosius Dobzhansky, celebrated this achievement by declaring that “nothing in biology makes sense except in the light of evolution” (Dobzhansky 1973).

As evolution was being recognized as the tie that binds the life sciences, a series of key papers was also launching the molecular revolution in biology. Crick articulated the central dogma of molecular biology in 1970, Danna and Nathans’ work on restriction endonucleases launched genetic engineering in 1971, and Sanger and coworkers published the dideoxy method for sequencing DNA in 1977. These advances opened the black box of Mendelian genetics, allowing biologists to unpack the molecular basis of heritable variation and differential reproductive success.

Researchers across the life sciences have embraced this Molecular Synthesis (Dean and Thornton 2007), and selection thinking is now pervasive among biologists working at scales ranging from protein-protein interactions to ecosystem dynamics. An avalanche of molecular data, combined with sophisticated, model-based methods in phylogeny inference (Felsenstein 2000), has also inspired an explosion in tree thinking, with applications ranging from BLAST searches (Camacho et al. 2009) to evo-devo research (Shubin et al. 2009, Tschopp and Tabin 2017) to evolutionary medicine and the study of emerging diseases (Grunspan et al. 2017).

How has the Molecular Synthesis impacted life sciences education? Although the seminal report on Vision and Change in Undergraduate Biology Education (AAAS 2011) listed evolution as its first core concept, researchers lament that many or most biology students are still being taught the modern synthesis’ version of evolution by natural selection. The concern is that instructors are missing the chance to use insights from molecular genetics if they do not ask students to make explicit connections between genotypes, phenotypes, and fitness (Hillis 2007, Smith et al. 2009, Kalinowski et al. 2010, Nehm and Ridgway 2011, Dauer et al. 2013, White et al. 2013, Walck-Shannon et al. 2019, Goransson et al. 2020).

Why has our teaching of natural selection lagged so far behind advances in research? One reason may be that evolution by natural selection, although “breathtakingly” simple in concept (Chown 2013), is notoriously difficult to teach (Nehm and Reilly 2007, Gregory 2009, Ha and Nehm 2013). For example, it requires students to adopt and apply perspectives that conflict with a series of ingrained cognitive biases (Nehm and Ridgway 2011, Coley and Tanner 2015, Richard et al. 2017). For example:

- *Population thinking* requires students to switch from a focus on the properties of individuals to the properties of populations (Gregory 2009). This requires them to abandon essentialism, which is the tendency to extend the attributes of individuals to a group (Coley and Tanner 2015). In evolutionary biology, one of essentialism’s consequences is a predilection to downplay variation among individuals. Accepting populations as the unit of evolution is also challenging because we routinely see the phenotypes of individuals change in response to environmental challenges in ways that look “adaptive,” making it easy to infer that these transformations cause evolutionary change. It is difficult for novices to parse that 1) acclimatized phenotypes are not passed on, but the ability to acclimatize is a heritable trait that responds to natural selection; and 2) natural selection acts on individuals, but only populations change.
- *Probabilistic thinking* is both non-intuitive and essential to thinking about evolutionary change (Nehm and Ridgway 2011, Goransson et al. 2020). It requires moving beyond counts to frequencies, and from predicting either/or outcomes to estimating likelihoods and rates of incremental change.
- *Incorporating a role for randomness and contingency and abandoning deterministic, goal-directed, and progressivist logic*. Research has shown that “everyday thinking” based on determinism, goal-directedness, or progressivism is the most common cause of naive ideas about natural selection (Goransson et al. 2020). For example, Darwin reminded himself to “Never use the words higher or lower” with respect to organisms, but today even professional biologists can slip into misleading references such as “higher plants” or “lower animals.” Recognizing the role of randomness and contingency also requires novices to overcome the human predilection to anthropomorphize and employ teleology (Coley and Tanner 2015). Too often, this task is made more difficult by the tendency of instructors to use teleological language as a shorthand when talking about adaptation (Gregory 2009).
- *Thinking across scales ranging from molecules to populations*. Recent research has emphasized the importance of this issue, which goes to the core of asking students to incorporate the Molecular Synthesis into their understanding of evolution by natural selection—specifically, to connect changes in genotypes to changes in phenotypes to changes in fitness to changes in the characteristics of populations (Bray Speth et al. 2009, Nehm and Ridgway 2011, Nehm and Ha 2011, Dauer et al. 2013, Bray Speth et al. 2014, Goransson et al. 2020). Humans have evolved to focus on visible phenomena and timescales of up to a few decades, but understanding evolution requires grasping events that are invisible and extremely fast, like DNA synthesis and repair, as well as historical events like the Cambrian “explosion,” which are largely invisible and occur over spans of tens of millions of years.
- *Synthesizing proximate and ultimate causation*. When students begin to explore the basics of cell biology and organismal biology, they face the challenge of not only recognizing the distinction between proximate (mechanistic) and ultimate (evolutionary) explanations for the phenomena they are studying, but also recognizing the two levels of explanation as complementary and then synthesizing them into a cohesive whole (Nehm and Ridgway 2011).
- *Socially constructed conflict around evolution.* For some students, evolution is a difficult topic because they regard it as a challenge to their personal religious beliefs (Barnes et al. 2020).

### Toward a solution: Using backward design to update our teaching approaches

Backward design is an approach to course structure that begins with articulating overall learning goals and specific learning objectives (Wiggins and McTighe 1998). Once these outcomes are established, instructors can align their teaching practices and their assessments to those outcomes, so that students can gain and demonstrate mastery of the learning objectives.

To teach evolution by natural selection in light of the Molecular Synthesis, it is essential to reconsider the learning objectives that were inspired by the Modern Synthesis. There are three key elements here:

1. Students should understand the contemporary definition of evolution as changes in allele frequencies, rather than the definition used by Darwin and the authors of the Modern Synthesis—the less-precise “changes in the characteristics of populations over time.” The explicit emphasis on alleles used by contemporary researchers is important because it makes the genetic basis of evolutionary change transparent.
2. Instructors should abandon the elaborate conceptual framework articulated by Darwin (1859) and the authors of the Modern Synthesis (Mayr 1982) in favor of explaining evolution by natural selection as an outcome of heritable variation associated with differential reproductive success (Nehm and Ha 2011, Ha and Nehm 2013, Bray Speth et al. 2014). The classical conceptual framework emphasized reproductive potential, resource limitation, population stability, and a “struggle for existence,” which are now considered to be unnecessary or even misleading ideas (Hillis 2007). For example, contemporary researchers recognize that natural selection acts in declining and expanding populations as well as stable ones, and via a wide array of processes and events—including disease, predation, mutualism, kin selection, sexual selection, and challenges posed by the physical environment—in addition to competition. Darwin and the authors of the Modern Synthesis also placed undue emphasis on survival as the key “currency” in evolution, even though differential reproductive success is the heart of evolutionary change (Gregory 2009).
3. Students should use the contemporary definition of mutation as a change in DNA sequence. They should also connect the physical mechanisms responsible for mutation to two key observations: mutation is random with respect to fitness, and it occurs in every population in every generation. Advanced undergraduates could be challenged to grasp that mutation rate itself is a heritable trait and can represent an adaptation to specific environments.

As an overall goal, instruction should focus on helping students connect changes in genotypes to changes in phenotypes to changes in fitness, and then recognize that evolution occurs as a logical outcome. To meet this goal, evolutionary biologists and education researchers have emphasized the importance of teaching the molecular basis of adaptation, using examples where the connections between genotype, phenotype, and fitness are understood — often at the level of the nucleotide (White et al. 2013, Reinagel and Bray Speth 2016).

The Molecular Synthesis provides an exciting framework for instructors to create learning objectives that reflect current research. Further, recent literature on teaching practice contains an increasing number of examples that instructors can use to help students understand the molecular nature of heritable variation and differential reproductive success (e.g. Kalinowski et al. 2010, White et al. 2013). Thus, instructors are equipped with a framework for writing learning objectives and exciting new examples to teach. The remaining challenge is to update how we assess student understanding of evolution by natural selection.

### A short history of evolution assessments

In addition to instructor-written questions that are used in formative and summative assessment, educators use two types of research-based, standardized tools to document novice-to-expert-level understanding of fundamental scientific ideas. One option is a concept inventory—a fixed-response, machine-gradable instrument that has distractors designed to reflect common but incorrect claims made by novices (Sands et al. 2018). A second option consists of open-response questions that are scored on a standardized rubric, which may or may not include points that are scaled to reflect gradations in understanding and/or deductions for declaring naïve ideas (e.g. Bishop and Anderson 1990). Rubrics provide a shared, research-based analytic framework for identifying and quantifying key elements in programs or writing (e.g. Wald et al. 2010, Chasteen and Scherr 2020) and are developed using an iterative process (Gehlbach and Brinkworth 2011).

Fixed-response concept inventories and open-response, rubric-based tools are grounded in literature that documents expert-level thinking and commonly-held naïve conceptions. They are assessed for validity and reliability (Moskal and Leydens 2000, Adams and Wieman 2010) and, once published, are administered before and after instruction for participation points only—so that completion, but not accuracy, affects the course grade. Their purpose is to provide instructors with diagnostic data that inform teaching practice or provide researchers with high-quality data to use in assessing the impact of experimental interventions.

To date, tools have been developed to assess student understanding of evolution by natural selection in both the concept inventory and rubric-based formats. The first published instruments were the Conceptual Inventory of Natural Selection, or CINS (Anderson et al. 2002), and what became known as the open-response instrument, or ORI (Bishop and Anderson 1990, Nehm and Reilly 2007). The CINS and ORI assessed expert understanding under the framework of the Modern Synthesis (Mayr 1982), and quantified common naive ideas about how natural selection works based on information in the literature (Bishop and Anderson 1990, Nehm and Reilly 2007). A more recent concept inventory, the Conceptual Assessment of Natural Selection, or CANS, placed an increased emphasis on the nature of mutation (Kalinowski et al. 2016), and the ORI was later developed into the Assessing Conceptual Reasoning of Natural Selection instrument, or ACORNS, by Nehm and co-workers (2012). The CINS and CANS both comprise questions about well-studied examples of adaptive change in morphology. In contrast, the ORI and ACORNS comprise variations on a framework that starts with a statement about a particular trait in a particular species and ends with a question of the form: “How would a biologist explain how [*exemplar trait*] in [*exemplar species*] evolved from an ancestor with [*ancestral trait*]?”

Nehm and co-workers not only developed a reliable computer-based scoring system for open responses on ACORNS questions (Ha et al. 2011, Moharreri et al. 2014) but also showed that the quality of student responses was context-dependent (Nehm and Ha 2011, Ha and Nehm 2013, see also Goransson et al. 2020). For example, the undergraduates they studied had more trouble explaining trait loss than trait gain, and the quality of students’ responses depended on whether they were tasked with explaining evolutionary change in 1) plants versus animals, or 2) familiar versus unfamiliar traits.

Although the CANS and ACORNS assessments are sophisticated, research-based, second-generation instruments, they are also grounded in part on concepts that were important historically, such as biotic potential, resource limitation, and competition or “struggle for existence.” They also do not emphasize the molecular nature of heritable variation and adaptation. To incorporate insights from the Molecular Synthesis more explicitly, we developed an expanded version of the ACORNS questions and scoring rubric that we call the E-ACORNS (Extended Assessing Conceptual Reasoning of Natural Selection) instrument.

### Developing the E-ACORNS

We had three primary goals in creating the E-ACORNS: 1) building on earlier advances by maintaining the ORI/ACORNS question framework; 2) challenging students to recognize the molecular basis of heritable variation and then make explicit connections between changes in genotypes, phenotypes, individual fitness, and allele frequencies in populations; and 3) updating the scoring rubric and increasing its resolution. We accomplished these goals by retaining the published ACORNS prompts but adding the sentence, “In your answer, be sure to connect what is happening at the molecular (genetic) level to the level of the whole organism,” and by creating a rubric that incorporated both insights from the Molecular Synthesis and gradations in scoring designed to capture changes in the sophistication of students’ thinking over time.

After an extensive literature search to review existing frameworks and assessments, our work on the new rubric continued by soliciting responses to E-ACORNS questions from experts in evolutionary biology, molecular biology, and biology education. Analyzing these responses allowed us to claim that an expert-level response would demonstrate understanding of five core concepts (Table 1):

1. Mutation is the source of heritable variation, and is random with respect to fitness;
2. Heritable variation in populations is due to variation in alleles, and exists irrespective of changes in the environment;
3. Changes in genotypes can lead to changes in phenotypes;
4. Selection occurs when certain phenotypes confer higher or lower fitness in a specific environment; and
5. Allele frequencies change due to fitness differences among alleles.

**Table 1.** The E-ACORNS rubric. Rubric elements that were considered molecular are identified with asterisks; other elements were considered nonmolecular.

| Core Concept | Novice | Intermediate | Advanced |
| --- | --- | --- | --- |
| **1. Nature of mutation** | *Mutation occurs, | *creates heritable variation, | *and is random with respect to fitness. |
| **2. Variation in populations** | Variation in populations exists, | *is based on a diversity of alleles, | *and exists independently of environmental conditions. |
| **3. Genotype to phenotype** | *Mutations change genotypes | *and may change gene products, | *and, if so, change phenotypes. |
| **4. Phenotype to fitness (natural selection)** | Traits vary in their impact on fitness, | leading to differential reproductive success | in a specific environment. |
| **5. Evolution** | Evolution occurs when trait frequencies change<br>— | or more precisely when allele frequencies change, | due to the fitness advantage of a trait. |

B. Naïve ideas
|  | 0 | -1 |
| --- | --- | --- |
| Teleological or anthropomorphic causation (purposeful or “conscious” change) | No mention | *Mutations occur in response to a change in the environment, or traits change due to want or need. |
| Inheritance of acquired characters | No mention | Traits change due to use/disuse, “exertion,” or interaction with the environment, with an implication that these changes are inherited. |
| Naïve group selectionism | No mention | Changes happen “for the good of the species.” |
| Essentialism | No mention | All individuals in a population change at once, or conflate adaptation with speciation. |

The first three concepts detail the molecular basis of heritable variation and its connection to phenotypic variation, responding to the call from Bray Speth and colleagues (2014, p. 529) for “assessment that promotes and reveals mechanistic and causal reasoning” about heritable variation. The fourth focuses on selection occurring via differential reproductive success, and the fifth on evolution as an outcome.

In addition to evaluating students’ understanding of natural selection, we also wanted to evaluate four naïve conceptions that are well-documented in the literature:

1. Teleological or anthropocentric explanations for the origin of either heritable variation or differential reproductive success (Bishop and Anderson 1990, Gregory 2009, Opfer et al. 2012, Richard et al. 2017);
2. Inheritance of acquired characters — meaning that characteristics developed during an organism’s lifetime are transmitted to offspring (Bishop and Anderson 1990, Gregory 2009, Ha et al. 2011);
3. “Good of the species” arguments based on naïve group selectionism (Anderson et al. 2002, Gregory 2009, Opfer et al. 2012); and
4. Essentialist claims focused on all individuals in a population changing in concert (Nehm and Reilly 2007, Gregory 2009, Richard et al. 2017).

We refined this rubric via extensive testing with introductory-level students. Specifically, we used student responses and the expert statements we had documented to propose gradations in mastery. For example, the second core concept in the rubric, which we summarized as “variation in populations,” is designed to indicate a novice level of understanding when students simply mention that traits vary within a population, an intermediate level when a response references a diversity of alleles in a population, and an expert-like level if students state that allelic diversity occurs irrespective of changes in the environment. This approach, which allows students’ increased sophistication to be measured over time, has been used in prior assessments focused on natural selection (Bishop and Anderson 1990, Opfer et al. 2012). Thus, the rubric for positive conceptions has a total of 15 elements: three levels for each of five major ideas. Naïve conceptions, in contrast, are scored on a simple presence-absence criterion, or 0-1, in the E-ACORNS rubric. Figure S1 illustrates the steps in drafting, revising, and testing the rubric.

The full rubric is given in Table 1. Table S1 provides a version with sample student responses to illustrate scoring decisions, and Table S2 compares elements in the E-ACORNS rubric with other instruments in the literature.

It is important to note that the E-ACORNS rubric is aspirational in nature — it was developed to reflect the ideas that a committee of experts would use to answer one of the prompts. By design, it is also heavily weighted toward insights from the Molecular Synthesis and the most challenging cognitive tasks for students: understanding the molecular basis of heritable variation and connecting changes in genotypes to changes in phenotypes and fitness (Nehm and Ridgway 2011, Dauer et al. 2013, Bray Speth et al. 2014, Reinagel and Bray Speth 2016, Walk-Shannon et al. 2019).

### A Practical Tool

Concept inventories and rubric-based assessments are useful because they quantify students’ understanding of fundamental concepts in biology. Instructors can use data from these research-based instruments in formative evaluation of teaching and respond by making changes in instructional practice; researchers can use the data as an outcome variable by analyzing students’ answers before and after an experimental intervention. Computerized scoring of the ACORNS rubric makes these applications simple for instructors (Ha et al. 2011, Moharreri et al. 2014). But is the E-ACORNS—which currently must be scored by hand—feasible to use as an instructional and research tool, especially at scale?

Our research offers three points relevant to answering this question. First, we achieved reliable scoring—assessed as interrater reliabilities calculated as Cohen’s Kappa that exceeded 0.80 or even 0.90—when advanced undergraduates who had been trained on the rubric for 5-6 hours served as raters. Once they had achieved this level of reliability, raters were able to work independently—meaning that we did not have to require two raters to evaluate each response and meet to reach consensus. Instead, we only had to schedule intermittent check-ins with the rating team to discuss their scores on a small sample of common questions. The goal of this step was to ensure that coder drift was not occurring. Once trained, raters were able to score a student response in less than two minutes, on average, meaning that a single undergraduate researcher could reliably evaluate over a hundred responses in an hour.

To test the utility of the E-ACORNS more directly, we conducted a preliminary study by administering the assessment to students in a multi-term introductory course for Biology majors. The initial term introduced evolution, Mendelian genetics, and ecology; the second term introduced biological molecules, cell biology, and molecular genetics. Although this ordering of topics is opposite the pattern at most institutions in North America, a clear separation between one term that emphasizes molecular biology and one term that focuses on evolution and whole-organism biology is almost universal in introductory course sequences. We used different prompts to assess student understanding at three time points over the two terms at our institution: at the start of the initial term, at the end of the initial term, and at the end of the second term. At each time point, students were presented with two prompts: one that asked them to explain a trait gain, and another that asked them to explain a trait loss (see Supplementary Materials for each time point’s specific prompts). We had raters score each of the 15 elements in the rubric as 1/0, and summed points for each student to get a total score. To obtain the maximum score of “15,” a response must demonstrate the full spectrum of understanding across all 3 columns of Table 1 for each of the five core concepts. We analyzed E-ACORNS scores from 327 students who completed the trait gain prompts on the first two assessments (317 for trait loss prompts) and 214 students who completed the trait gain prompts on all three assessments (196 for Loss).

After two terms of introductory biology, the median E-ACORNS scores for students at our institution were 3 points (for trait-gain prompts) and 2 points (for trait-loss prompts) out of 15 points possible. These findings are consistent with the cognitive challenges involved in understanding evolution by natural selection noted earlier, and empirical research suggesting that less than 10% of undergraduates have a “functional understanding” of the process (Gregory 2009, p. 163). An examination of percent correct for each element in the rubric (Table S3) suggested that student scores after the second term conformed to the novice > intermediate > expert-like sequence proposed in the rubric (Table 1), although that pattern was rarely observed in scores analyzed after the first term. In addition, scores after the second course appeared to be exceptionally low on items related to 1) variation in populations, and 2) the link between fitness and changes in allele frequencies.

To assess patterns in student performance in more detail, we explored whether student scores improved over the two terms and whether responses differed on molecular versus nonmolecular components of the rubric. Regression models showed that overall scores *declined* on both trait gain and trait loss prompts over the two terms of the introductory series: Scores when students started the introductory series for majors were higher than scores after six months of instruction (Figure S2, Table S4). This result occurred even though, on both trait gain and trait loss prompts, students did better on molecular versus nonmolecular components of the rubric in the second term (Figure S3, Table S5, Table S6) — suggesting that scores on nonmolecular components had a disproportionately large decline in the second term. Part of the overall drop in scores could be explained by the prompt that was administered after the second term being more difficult, as it was based on unfamiliar taxa and traits, but the data also indicated no increase in scores on the identical prompts, focused on familiar taxa and traits, used at the start and end of the first course (Table S4). Our data do indicate, however, that students declared many fewer naïve ideas at the end of the second course in response to both trait gain or trait loss prompts (Table S4C).

In contrast to the patterns observed with responses to the ACORNS instrument, where answers have been found to be highly context-dependent (see Nehm & Ha 2011), we found no significant differences between scores on trait-gain versus trait-loss questions (Table S7). We do not know if our result is due to the increased emphasis on molecular mechanisms in the E-ACORNS rubric, an unusual sample of students at our institution, or other causes. If the same lack of contrast were observed in other student populations, it would suggest that instructors and researchers can simplify the task of assessing students’ understanding of natural selection by using either a trait-gain or trait-loss prompt, instead of both.

Taken together, our preliminary study suggests that this introductory biology course sequence did not support robust gains in students’ understanding of evolution by natural selection, even though an encouraging reduction in naïve conceptions occurred. To explain why, we hypothesize that students’ responses reflected the siloed nature of instruction. Specifically, students in the initial course got a solid grounding in the Modern Synthesis’ version of evolution by natural selection, supplemented by a single class-session that introduced the molecular basis of mutation. In this way, instruction in the first term focused on the connection between phenotypes and fitness. Students in the second course, in contrast, got a solid grounding in the Central Dogma and molecular genetics, and thus the connection between genotypes and phenotypes. The second course focuses on canonical examples of molecular processes, however, and ignores intraspecific and interspecific variation. This lack of emphasis on variation may explain the exceptionally low scores we observed for this section of the E-ACORNS rubric. Further, the genotype-phenotype connection in the second course is taught with almost no reference to fitness and evolution by natural selection. In this introductory biology series, as in most curricula, students were never asked to make explicit connections between genotypes, phenotypes, fitness, and changes in allele frequencies in populations. The E-ACORNS assessment revealed that in these students’ minds, the molecular synthesis had not yet occurred.

## Conclusions and future directions

The E-ACORNS assessment provides an effective and efficient diagnostic tool for instructors who want to assess their students’ understanding of natural selection in more depth than is possible with typical quiz or exam questions. It also offers a new instrument for researchers to quantify outcomes on teaching innovations focused on evolution. Future work could start with efforts to test the rubric in additional student populations, to assess the generality of the patterns presented here. Further research could modify the rubric based on additional expert opinion and, ideally, work that documents the learning progressions (see Duschl 2019) involved in the five core concepts articulated here, to see if the steps that students go through as they transition from novice- to intermediate- to expert-like understanding conform to the steps proposed in the current E-ACORNS rubric. It would also be helpful to administer E-ACORNS to upper division undergraduates and beginning graduate students, to assess whether their overall performance is better than beginning undergraduates’ and whether their responses conform to the intermediate and expert-like levels of understanding reflected in Table 1.

Given the current structure of most introductory biology courses for majors, however, it may be difficult for faculty to document meaningful gains in how well their first-year students understand evolution by natural selection in terms of the Molecular Synthesis. Currently, the initial semester of most year-long courses focuses on biomolecules, cell biology, and molecular genetics, with the second semester introducing evolution, diversity of life, physiology, and ecology. The major textbooks follow the same sequence. Because the teaching teams for the two semesters are almost always distinct—sometimes even being housed in different departments at the same institution—it is understandable that students have trouble integrating levels of organization and grasping the Molecular Synthesis. Our preliminary data appear to document this difficulty.

To remedy the situation, we urge faculty to consider starting introductory courses for majors and non-majors by introducing the central dogma of molecular biology and its connection to evolution by natural selection, using well-researched examples that are still under investigation (Smith et al. 2009, White et al. 2014). In courses for majors, subsequent material on biomolecules, cell structure and function, and the details of molecular genetics should be even more meaningful. In nonmajors courses, subsequent material on issues of human interest can be scaffolded on a solid conceptual foundation. In both cases, instruction will do a better job of reflecting the nature of contemporary work in bioscience, where traditional boundaries between subdisciplines are thinning, and researchers are increasingly fluent in the languages of both molecular and evolutionary biology.

## Acknowledgments

We thank Elizabeth Glenski, Grace E.C. Dy, Mariah Hill, and Elisa Tran for assistance with analyzing expert responses to E-ACORNS questions. This work was made possible by a grant from the Howard Hughes Medical Institute, number 52008126, and was conducted under University of Washington Human Subjects Division application 00003631.

## Notes

Conflict of interest: The authors declare no conflicts of interest, including conflicts based on employment affiliation or financial interests.

### Competing Interest Statement

The authors have declared no competing interest.

## References cited

Adams WK, Wieman CE. 2010. Development and validation of instruments to measure learning of expert-like thinking. International Journal of Science Education doi.org/10.1080/09500693.2010.512369.

AAAS (2011). Vision and Change in Undergraduate Biology Education: A Call to Action. Washington, DC: AAAS. Available online at http://visionandchange.org/finalreport/.

Anderson DL, Fisher KL, Norman GJ. 2002. Development and evaluation of the conceptual inventory of natural selection. Journal of Research on Science Teaching 39: 952–978.

Barnes ME, Dunlop HM, Sinatra GM, Hendrix TJ, Zheng Y, Brownell SE. 2020. “Accepting evolution means you can’t believe in God”: atheistic perceptions of evolution among college biology students. CBE-Life Sciences Education 19: ar21, 1–13.

Bishop BA, Anderson CW. 1990. Student conceptions of natural selection and its role in evolution. Journal of Research on Science Teaching 27: 415–427.

Bray Speth E, Long TM, Pennock RT, Ebert-May D. 2009. Using Avida-ED for teaching and learning about evolution in undergraduate introductory biology courses. Evolution: Education and Outreach 2: 415–428.

Bray Speth E, Shaw N, Momsen J, Reinagel A, Le P, Taqieddin R, Long T. 2014. Introductory biology students’ conceptual models and explanations of the origin of variation. CBE-Life Sciences Education 13: 529–539.

Brownell SE, Freeman S, Wenderoth MP, Crowe AJ. 2014. BioCore Guide: a tool for interpreting the core concepts of Vision and Change for Biology majors. CBE-Life Sciences Education 13: 200–211.

Camacho C, Coulouris G, Avagyan V, Ma N, Papadoopoulos J, Bealer K, Madden TL. 2009. BLAST+; architecture and applications. BMC Bioinformatics 10: 421–430.

Chasteen SV, Scherr RE. 2020. Developing the physics teaching education program analysis rubric: measuring features of thriving programs. Physical Review Physics Education Research 16, 010115.

Chown M. 2013. What a Wonderful World. Faber and Faber.

Coley JD, Tanner K. 2015. Relations between intuitive biological thinking and biological misconceptions in biology majors and nonmajors. CBE-Life Sciences Education 14: 1–19.

Crick F.. 1970. Central dogma of molecular biology. Nature 227: 561–563.

Danna K, Nathans D. 1971. Specific cleavage of simian virus 40 DNA by restriction endonuclease of *Hemophilus influenzae*. Proceedings of the National Academy of the USA 68: 2913–2917.

Darwin, C. 1859. The Origin of Species. Simon & Schuster.

Dauer JT, Momsen JL, Bray Speth, E, Makohon-Moore SC, Long TM. 2013. Analyzing change in students’ gene-to-evolution models in college-level introductory biology. Journal of Research on Science Teaching 50: 639–659.

Dean AM, Thornton JW. 2007. Mechanistic approaches to the study of evolution: the functional synthesis. Nature Reviews Genetics 8: 675–688.

Dobzhansky T. 1973. Nothing in Biology makes sense except in the light of evolution. The American Biology Teacher 35: 125–129.

Duschl RA. 2019. Learning progressions: framing and designing coherent learning sequences for STEM education. Disciplinary and Interdisciplinary Science Education Research doi.org/10.1186/s43031-019-0005-x.

Felsenstein J. 2004. Inferring Phylogenies. Sinauer.

Gehlbach H, Brinkworth ME. 2011. Measure twice, cut down error. Harvard Library https://dash.harvard.edu/handle/1/8138346.

Goransson A, Orraryd D, Fiedler D, Tibell LAE. 2020. Conceptual characterization of threshold concepts in student explanations of evolution by natural selection and effects of item context. CBE-Life Sciences Education 19: ar1.

Gregory TR. 2009. Understanding natural selection: essential concepts and common misconceptions. Evolution: Education and Outreach 2: 156–175.

Grunspan DZ, Nesse RM, Barnes ME, Brownell SE. 2017. Core principles of evolutionary medicine. Evolution, Medicine, and Public Health 2018: 13–23.

Ha M, Nehm RH, Urban-Lurain M, Merrill JE. 2011. Applying computerized-scoring models of written biological explanations across courses and colleges: prospects and limitations. CBE-Life Sciences Education 10: 379–393.

Ha M, Nehm RH. 2013. Darwin’s difficulties and students’ struggles witih trait loss: cognitive-historical parallelisms in evolutionary explanation. Science & Education doi 10.1007/s11191-013-9626-1.

Hillis DM. 2007. Making evolution relevant and exciting to biology students. Evolution 61: 1261–1264.

Kalinowski ST, Leonard, MJ, Andrews TM. 2010. Nothing in evolution makes sense except in the light of DNA. CBE-Life Sciences Education 9: 87–97.

Kalinowski ST, Leonard MJ, Taper ML. 2016. Development and validation of the Conceptual Assessment of Natural Selection (CANS). CBE-Life Sciences Education 15:ar64, 1.

Mayr E. 1982. The Growth of Biological Thought. Harvard University Press.

Moharreri K, Ha M, Nehm RH. 2014. EvoGrader: an online formative assessment tool for automatically evaluating written evolutionary explanations. Evolution: Education and Outreach 7: 15–28.

Moskal BM, Leydens JA. 2000. Scoring rubric development: validity and reliability. Practical Assessment, Research & Evaluation. 7: 1–6.

Nehm RH, Beggrow EP, Opfer JE, Ha M. 2012. Reasoning about natural selection: diagnosing contextual competency using the ACORNS instrument. The American Biology Teacher 74: 92–98.

Nehm RH, Ha M. 2011. Item feature effects in evolution assessment. Journal of Research in Science Teaching doi 10.1002/tea.20400.

Nehm RH, Reilley L. 2007. Biology majors’ knowledge and misconceptions of natural selection. BioScience 57: 263–272.

Nehm RH, Ridgway J. 2011. What do experts and novices “see” in evolutionary problems? Evolution: Education and Outreach 4: 666–679.

Opfer JE, Nehm RH, Ha M. 2012. Cognitive foundations for science assessment design: knowing what students know about evolution. Journal of Research in Science Teaching 49: 744–777.

Reinagel A, Bray Speth E. 2016. Beyond the central dogma: model-based learning of how genes determine phenotypes. CBE-Life Sciences Education 15: 1–13.

Richard M, Coley JD, Tanner KD. 2017. Investigating undergraduate students’ use of intuitive reasoning and evolutionary knowledge in explanations of antibiotic resistance. CBE-Life Sciences Education 16: ar55.

Sands D, Galloway RK, Jordan S. 2018. Using concept inventories to measure understanding. Higher Education Pedagogies 3: 60–69.

Sanger F, Nicklen S, Coulson AR. 1977. DNA sequencing with chain-terminating inhibitors. Proceedings of the National Academy of the USA 74: 5463–5467.

Shubin N, Tabin C, Carroll S. 2009. Deep homology and the origins of evolutionary novelty. Nature 457: 818–823.

Smith JJ, Baum DA, Moore A. 2009. The need for molecular genetic perspectives in evolutionary education (and vice versa). Trends in Genetics 25: 427–429.

Tschopp P, Tabin CJ. 2017. Deep homology in the age of next-generation sequencing. Philosophical Transactions of the Royal Society of London B 372: 20150475.

Walck-Shannon E, Batzli J, Pultorak J, Boehmer H. 2019. Biological variation as a threshold concept: can we measure threshold crossing? CBE-Life Sciences Education 18: ar36.

Wald HS, Borkan JM, Scott Taylor M, Anthony D, Reis SP. 2010. Fostering and evaluating reflective capacity in medical education: developing the REFLECT rubric for assessing reflective writing. Academic Medicine 87: 41–50.

White PJT, Heidemann M, Loh M, Smith JJ. 2013. Integrative cases for teaching evolution. Evolution: Education and Outreach 6: 17–23.

Wiggins G, McTighe J. 1998. Understanding by Design. Association for Supervision and Curriculum Development.

