## Supplementary material for "From the Modern Synthesis to the Molecular Synthesis: updating how we teach and assess evolution by natural selection": Supplementary Materials.docx

I. **Rubric development**

**Figure S1 Steps in developing the E-ACORNS rubric**

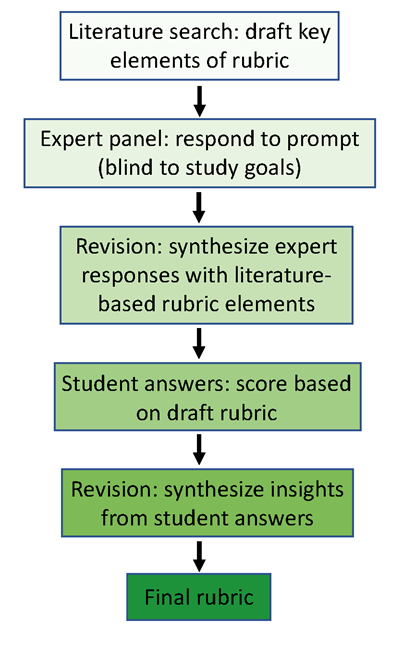

**Table S1 E-ACORNS rubric with sample student responses**

The student responses provided here illustrate how raters connected actual student responses to specific elements the rubric. Words or phrases that were key to scoring decisions for the specific concept listed in each row are italicized.

| **Expert-like idea** | **Relevant portion of example student response** |
| --- | --- |
| 1. Nature of mutation | |
| Mutation occurs, | A *mutation occurred*. |
| creates heritable variation, | Over time, being flightless benefited birds and those individuals mated and *passed on their traits* onto their offspring. |
| and is random with respect to fitness. | The gene then *randomly mutated* and turned into something that was venom-like. |
| 2. Variation in populations | |
| Variation in populations exists, | The *genes that code for the trait* of poison production *have heritable variation* that can be passed on to offspring. |
| is based on a diversity of alleles, | There was *genetic variation in the ancestral species of snail from mutations*, such as some snails would be poisonous and others are not. |
| and exists independently of environmental conditions. | Within the population of non-poisonous snails in the past, there was genetic variation. *Due to the genetic variation caused by a mutation, crossing over, etc., there were snails who were poisonous. Throughout time, pressures were applied* to the whole snail population - whether it be fragmentations in the environment or natural selection, poisonous snails had higher fitness. With their higher fitness, they had higher reproductive success and are now considered descendants of snails that were not poisonous. |
| 3. Genotype to phenotype | |
| Mutations change genotypes, | From a population of non-poisonous snails, *a mutation occurred* within their gene that led to *an allele for the poisonous trait*. |
| and may change gene products, | At the molecular level, the DNA in the organism has coding that either has been changed through some sort of *mutation* (point/ deletion/ shift) and *caused a change in phenotype* or if the gene was not being expressed in the ancestors but now is expressed. |
| and, if so, change phenotypes. | Probably, there would be a few mutations in the DNA sequence that lead to different amino acid products, *changing the protein products and leading to a different characteristic* of that organism |
| 4. Phenotype to fitness | |
| Traits vary in their impact on fitness, | Since that trait would be beneficial to the fitness of said snail, the snail's *fitness would increase compared to the rest* of the species, increasing the number of offspring from the one snail. |
| leading to differential reproductive success | Since that trait would be beneficial to the fitness of said snail, the snail's fitness would increase compared to the rest of the species, *increasing the number of offspring from the one snail.* |
| in a specific environment. | Individuals with the ability to produce poison had *a fitness advantage in one environment*, and therefore higher reproductive success. |
| 5. Evolution | |
| Evolution occurs when trait frequencies change— | Individuals with the ability to produce poison had a fitness advantage in one environment, and therefore higher reproductive success. Over time, these alleles increased in frequency in the population, which *led to the entire population of snails having this trait*. |
| or more precisely when allele frequencies change, | Individuals with the ability to produce poison had a fitness advantage in one environment, and therefore higher reproductive success. Over time, *these alleles increased in frequency* in the population, which led to the entire population of snails having this trait. |
| due to the fitness advantage of a trait. | Individuals with the ability to produce poison *had a fitness advantage* in one environment, and *therefore higher reproductive success*. Over time, these alleles increased in frequency in the population, which led to the entire population of snails having this trait. |
| **Naive idea** | **Relevant portion of example student response** |
| 1. Teleological or anthropomorphic causation | The snail developed the poison because *they needed* to protect itself. |
| 2. Inheritance of acquired characters | Other explanations that biologists could potentially use would be the surfacing of this evolutionary toxicity as a result of a defensive needs. The *gradual change in the snail's surroundings and natural predators and competition could spur the development of its poisonous trait* in order for the snail to defend itself and deter predators |
| 3. Naïve group selectionism | The new species of snail is poisonous because the environment demands the change to occur *for the survival of the species*. In the ancestral snails did not require the need of protecting themselves from predators - if they had means to defend themselves, it no longer works for the new species. When the pregnant ancestry snail could not defend against itself from predators, its genetic coding changed so that the offspring had a better chance of surviving. |
| 4. Essentialism | As the population of snails continued to grow, *more and more snails become poisonous, creating a new species*. |

**Table S2 Concepts included in instruments used to assess student understanding of natural selection**

In each case, a member of the research team (SF) scored statements about each concept from data in tables, figures, or text in the citations noted. “X” denotes that the instrument contains that concept.

A.	Expert-like ideas

| **Concept** | **Instrument** | | | | | |
| --- | --- | --- | --- | --- | --- | --- |
|  | CINS: Anderson et al. 2002 | ORI: Nehm & Reilly 2007 | ACORNS: Ha et al. 2011 | EvoGrader: Moharreri et al. 2014 | CANS: Kalinowski et al. 2016 | E-ACORNS |
| Biotic potential | X | X |  |  | X |  |
| Population stability | X |  |  |  |  |  |
| Resource limitation | X | X | X | X |  |  |
| Limited survival | X | X | X | X | X |  |
| Variation in populations | X |  | X | X | X | X |
| Heritable variation | X | X | X | X | X | X |
| Sources of variation | X | X |  |  | X | X |
| Differential survival | X | X | X | X | X |  |
| Differential reproductive success |  |  |  |  |  | X |
| Evolution as outcome | X | X |  |  | X | X |
| Origin of species | X |  |  |  |  |  |
| Role of chance in success |  |  |  |  | X |  |
| Genotype to phenotype |  |  |  |  | X | X |
| Role of environmental stress in success |  |  |  |  | X |  |

Table S2, continued

B. Naïve ideas

| **Concept** | **Instrument** | | | | | | |
| --- | --- | --- | --- | --- | --- | --- | --- |
|  | Bishop & Anderson 1990 | CINS: Anderson et al. 2002 | ORI: Nehm & Reilly 2007 | ACORNS: Ha et al. 2011 | EvoGrader: Moharreri et al. 2014 | CANS: Kalinowski et al. 2016 | E-ACORNS |
| Goal-directed change (teleology) | X | X | X |  | X | X | X |
| Change via use/disuse | X |  | X |  | X | X | X |
| Change in individuals is transmitted | X | X |  |  | X |  | X |
| Species change as a unit | X | X | X |  |  |  | X |
| Traits themselves change | X |  |  |  |  |  |  |
| Change for the good of the species |  | X |  |  |  |  | X |
| Change via intention (anthropomorphic) |  | X |  |  |  |  |  |

II. **Data analysis from preliminary study of student responses to trait-gain and trait-loss prompts**

We included E-ACORNS prompts in a large survey that was conducted for a different study and administered to students in the introductory biology series for majors at the University of Washington (Mukerji et al. in prep). The first term in this 3-quarter series, Biology 180 (hereafter “Bio1”), covers evolution, Mendelian genetics, diversity of life, and ecology, with one class period devoted to introducing the central dogma and molecular basis of mutation. The second course in the series, Biology 200 (hereafter “Bio2”), covers biological molecules, cell biology, molecular genetics, and animal development. The courses are taken primarily by 1st- and 2nd-year students; majors and non-majors take Bio1, while typically only majors proceed to Bio2. During the study, students were given participation points for completing the entire survey each time it was offered. The assignment included the following prompts:

At the start of Bio1 and end of Bio1,

- Trait Gain

“A species of snail (an animal) is poisonous. How would biologists explain how this species evolved from an ancestral species of snail that was not poisonous?

In your answer, be sure to connect what is happening at the molecular (genetic) level to the level of the whole organism.”

- Trait Loss

“A species of flightless bird (flightless birds, such as penguins, cannot fly) is closely related to bird species that are able to fly. How would biologists explain how a flightless bird species evolved from an ancestral bird species that was able to fly?

In your answer, be sure to connect what is happening at the molecular (genetic) level to the level of the whole organism.”

At the end of Bio2:

- Trait Gain

“A species of prosimian (an animal) has long tarsi. How would biologists explain how long tarsi evolved from an ancestral species of prosimian with short tarsi?

In your answer, be sure to connect what is happening at the molecular (genetic) level to the level of the whole organism.”

- Trait Loss

“A species of Suricata (an animal) with no pollex is closely related to species of Suricata that have a pollex. How would biologists explain how Suricata species that lack a pollex evolved from Suricata species that had a pollex?

In your answer, be sure to connect what is happening at the molecular (genetic) level to the level of the whole organism.”

After Bio2 concluded, we archived student responses to the E-ACORNS prompts and trained three advanced undergraduate students to score them on the rubric we had developed (see Table 1 and Table S1). Raters were paid. They were blind to the nature and purpose of the study, the identity of the participating students, and whether responses were from Pre-Bio1, Post-Bio1, or Post-Bio2. After a brief training on the rubric and a group discussion of how to score a small sample of student answers, a member of the research team conducted a series of more extensive training and norming sessions. During these, two or more raters independently scored the same sample of 10-20 responses, followed by the research team member facilitating a group discussion where responses and ratings were compared, contrasted, and discussed until raters reached consensus. This process was repeated in the initial meeting and in at least one additional meeting until inter-rater reliabilities on independently scored responses, calculated as Cohen’s Kappa, were above 0.80. This is the “high” standard used in the evaluation literature as a benchmark, above which raters can work independently (Koo and Li 2017). Once raters had qualified in this way, they were given different responses to evaluate on their own, in batches of 100-200. Each time they worked through 2-3 of these batches, a member of the research team facilitated a meeting designed to guard against “coder drift” by computing inter-rater reliabilities and comparing, contrasting, and discussing the judgments they had made on groups of 10-20 common responses.

Raters scored each of the 15 points in the rubric as present (1) or absent (0), and the four naive conceptions as present (-1) or absent (0). We then used the sum of total correct conceptions and the sum of total naive ideas as the independent variables in separate regression models and other analyses focused on the following four research questions:

1. Which elements of the rubric are easier or more difficult than others?
2. Did responses improve over time, as students progressed through the two 10-week terms?
3. Did performance on the molecular concepts (items marked “*” in Table 1) differ from performance on nonmolecular concepts?
4. Did performance on prompts about trait gains differ from performance on prompts about trait loss?

We answered Research Question 2 (RQ 2) with paired two sample t-tests, and RQs 3-4 by fitting multilevel binomial regression models, modeling the outcome as the proportion of the points earned on the rubric. We used the Akaike Information Criterion (AIC; Anderson and Burnham 2004) to confirm that binomial models, which treat the outcome as a proportion of the total possible score, fit better than either proportional odds logistic regression models, which treat the outcome as an ordinal categorical variable, or linear models (see Theobald et al. 2019). For RQ 3, we included an indicator for whether the rubric item was a molecular or nonmolecular item. For RQ 4, we included a factor indicating whether the score came from the gain or loss question. The basic models were as follows:

1. PostBio1 ~ PreBio1 + indicator
2. PostBio2 ~ PreBio1 + indicator
3. PostBio2 ~ PostBio1 + indicator
4. PostBio2 ~ PreBio1 + PostBio1 + indicator
5. Post Bio2 ~ PreBio1*Indicator + PostBio1*Indicator

Further, each model included a random intercept for each student, acknowledging that in this repeated-measures design, responses from individual students are not independent (Theobald 2018), and that for each student, responses to questions about trait gains are not wholly independent from responses to questions about trait losses.

Together, the models test that, all else equal, the odds of earning more points on molecular aspects of the rubric are the same as nonmolecular aspects and that the odds of earning more points when describing trait gain are the same as describing trait loss. All analyses were performed in R Version 4.0.3 (R Core Team 2021).

***Reasearch Question 1 (RQ 1): Which elements of the rubric are easier or more difficult than others?***

Because the E-ACORNS rubric is newly developed, we generated descriptive data by computing the percent correct in a large sample of student responses for each of the 15 elements in the positive conceptions rubric and the percent declared for each of the four items in the naive ideas rubric. Results for responses at the start and end of the initial course and at the end of the second course are reported in Table S3.

**Table S3 Percent of trait-gain prompt responses that included individual components of the E-ACORNS rubric**

Numbers in parentheses are sample sizes; asterisks (*) indicate molecular concepts.

A. Expert-like ideas

| **Concept** | **PreBio1 (380)** | **PostBio1 (327)** | **PostBio2 (214)** |
| --- | --- | --- | --- |
| Nature of mutation |  |  |  |
| *Mutation occurs, | 50.5 | 60.9 | 53.7 |
| *creates heritable variation, | 25.8 | 22.6 | 28.0 |
| *and is random with respect to fitness. | 27.1 | 40.4 | 11.2 |
| Variation in populations |  |  |  |
| Variation in populations exists, | 50.0 | 48.0 | 17.3 |
| *is based on a diversity of alleles, | 3.4 | 21.1 | 6.1 |
| *and exists independently of environmental conditions. | 36.3 | 39.4 | 0.0 |
| Genotype to phenotype |  |  |  |
| *Mutations change genotypes, | 39.7 | 31.2 | 27.6 |
| *and may change gene products, | 56.3 | 39.4 | 20.6 |
| *thus changing phenotypes. | 2.9 | 0.3 | 6.1 |
| Phenotype to fitness |  |  |  |
| Traits vary in their impact on fitness, | 77.6 | 74.3 | 65.4 |
| leading to differential reproductive success | 38.7 | 24.8 | 17.8 |
| in a specific environment. | 55.0 | 39.4 | 10.3 |
| Evolution |  |  |  |
| Evolution occurs when trait frequencies change— | 61.6 | 44.0 | 21.5 |
| or more precisely when allele frequencies change— | 3.4 | 24.8 | 3.7 |
| due to the fitness advantage of a trait. | 2.9 | 22.3 | 0.0 |

B. Naïve ideas

| **Naive idea** | **PreBio1 (380)** | **PostBio1 (327)** | **PostBio2 (214)** |
| --- | --- | --- | --- |
| Teleological or anthropomorphic causation | 27.6 | 14.7 | 0 |
| Inheritance of acquired characters | 2.9 | 3.1 | 1.4 |
| Naïve group selectionism | 29.2 | 36.7 | 0 |
| Essentialism | 13.7 | 15.0 | 0.9 |

***Research Question 2: Did responses improve over time, as students progressed through the two 10-week terms?***

Because some of the concepts in the E-ACORNS rubric are covered in the initial course while others are a focus of the second course, we predicted that student performance on both the trait gain and trait loss prompts would improve over time. To test this prediction, we performed Welch’s paired *t*-tests to compare student responses across terms to both trait gain and trait loss prompts.

As the results reported in Table S4A indicate, there was no change in student scores on trait-gain prompts from the start to the end of the initial course. There was, however, a statistically significant drop in average scores from the end of the initial course to the end of second course, as well as a drop from the before the initial course to the end of the second course. These results indicate that instead of improving over time, student performance on trait-gain questions declined from the start of the initial term to the end of the second term, and that much or most of the drop occurred in the second course, where evolution is not a prominent part of the topic context or coverage. The violin plots in Figure S2A visualize this result by indicating the medians, the interquartile ranges, and a kernel density representation of the data for trait gain responses in Pre-Bio1, Post-Bio1, and Post-Bio2.

**Table S4 Results of paired *t-*tests for student responses across time**

A. Expert-like ideas: Trait gain

|  | Mean Difference | *t*-value | df | *p*-value |
| --- | --- | --- | --- | --- |
| PreBio1 vs. PostBio1 | -0.33 | -1.42 | 213 | 0.1571 |
| PostBio1 vs. PostBio2 | -2.03 | -9.59 | 213 | < 0.0001 |
| PreBio1 vs. PostBio2 | -2.36 | -10.42 | 213 | < 0.0001 |

B. Expert-like ideas: Trait loss

|  | Mean Difference | *t*-value | df | *p*-value |
| --- | --- | --- | --- | --- |
| PreBio1 vs. PostBio1 | -0.35 | -1.94 | 195 | 0.053 |
| PostBio1 vs. PostBio2 | 0.26 | 1.56 | 195 | 0.119 |
| PreBio1 vs. PostBio2 | -0.09 | -0.46 | 195 | 0.643 |

C. Naive ideas: Trait gain

|  | Mean Difference | *t*-value | df | *p*-value |
| --- | --- | --- | --- | --- |
| PreBio1 vs. PostBio1 | 0.01 | 0.13 | 213 | 0.90 |
| PostBio1 vs. PostBio2 | -0.59 | -10.36 | 213 | < 0.0001 |
| PreBio1 vs. PostBio2 | -0.58 | -10.39 | 213 | < 0.0001 |

D. Naive ideas: Trait loss

|  | Mean Difference | *t*-value | df | *p*-value |
| --- | --- | --- | --- | --- |
| PreBio1 vs. PostBio1 | -0.01 | -0.34 | 195 | 0.74 |
| PostBio1 vs. PostBio2 | -0.10 | -2.88 | 195 | 0.004 |
| PreBio1 vs. PostBio2 | -0.11 | -2.94 | 195 | 0.004 |

Results from analyzing responses on trait loss questions indicate that scores were stable over the three time points (Table S4B, Figure S2B). Thus, neither trait gain nor trait loss data support the hypothesis that student understanding of evolution by natural selection increased over time during this introductory course series for majors.

Evaluating changes in the number of naive ideas that students declared in trait gain or trait loss questions (Table S4C-D) is more challenging because, in general, students did not hold many naïve ideas. As Figure S2C shows, the small number of students who indicated naive ideas in the PreBio1 and PostBio1 responses dwindled over time, so that almost all values were zero for total naive ideas in the PostBio2 survey.

**Figure S2 Violin charts depicting E-ACORNS scores of student responses across two terms**

A. Positive conceptions

Trait gain prompts	Trait loss prompts

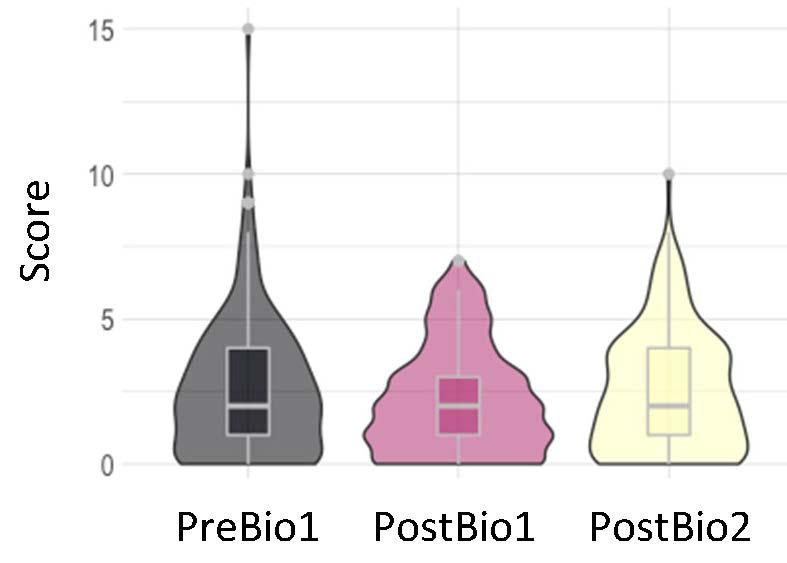

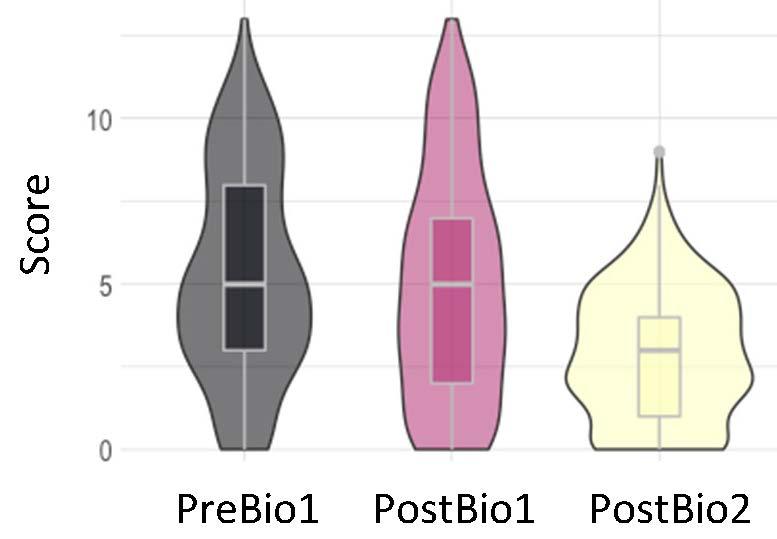

B. Naive ideas

Trait gain prompts	Trait loss prompts

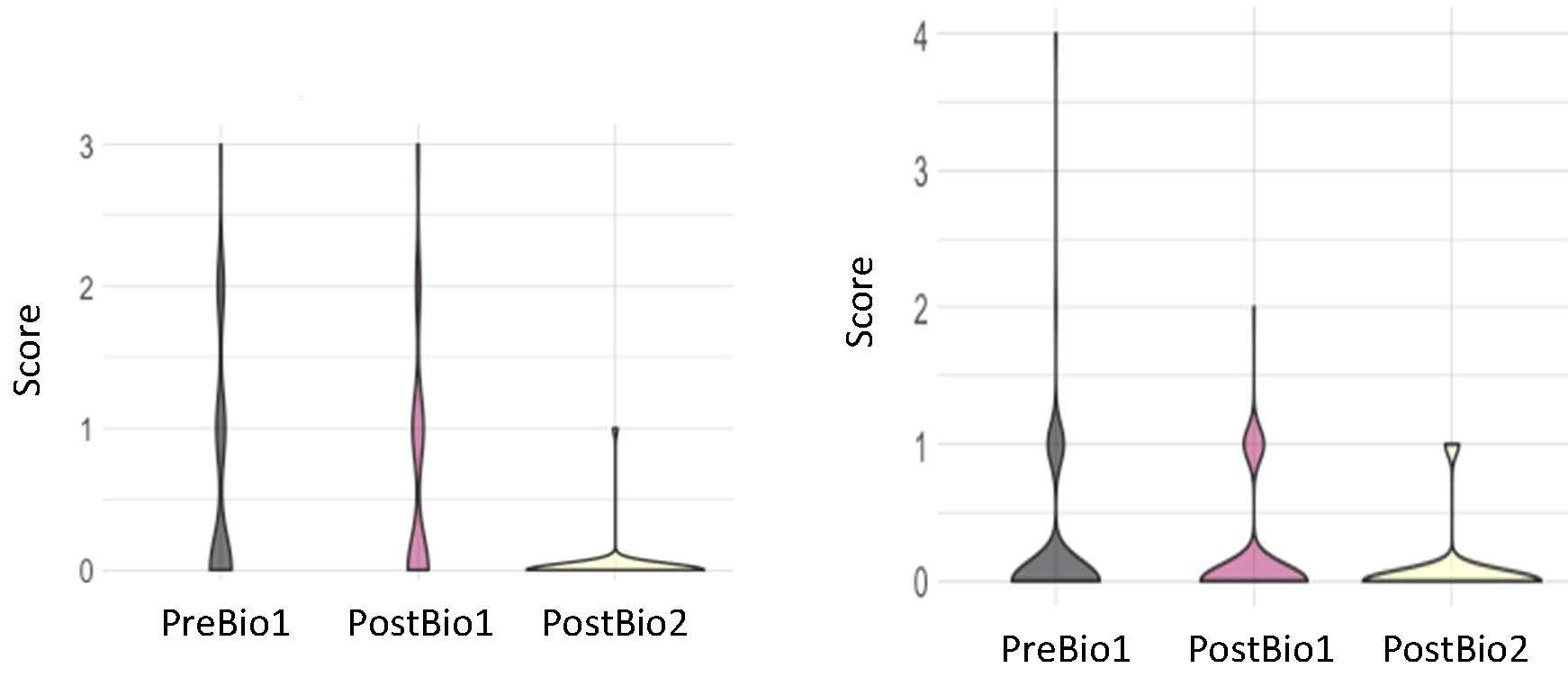

***Research Question 3: Did performance on molecular concepts differ from performance on nonmolecular concepts?***

This question was motivated by the E-ACORNS’ goal of quantifying how well students understand the molecular basis of evolutionary change, which research has shown is particularly difficult (Bray Speth et al. 2009, Dauer et al. 2013, Speth et al. 2014, Walck-Shannon et al. 2019, Goransson et al. 2020). The course context in this preliminary study was appropriate to address the question, as the initial course introduces students to the Modern Synthesis, with molecular material following in the second term. During instruction there was, however, no explicit effort to tie genotypes, phenotypes, and fitness together—a central goal of the Molecular Synthesis approach to teaching evolution by natural selection.

To explore RQ 3, we modeled post-course scores on trait gain responses as a function of one or more prior scores, a dummy variable indicating whether the rubric-item was nonmolecular or molecular as a fixed effect, and a random variable identifying individual students. The molecular items of the E-ACORNS rubric (see Table 1) comprised all three elements in each of the *Nature of Mutation* and *Genotype-to-phenotype* concepts, along with the “is based on a diversity of alleles” item in the *Variation in Populations* element. The other eight items in the rubric were coded as nonmolecular. Table S6 summarizes the models and regression output for trait gain questions, but note that we also proposed three models that turned out to have a singular fit:

- PostBio2 ~ PostBio1 + NonMolecular + (1|Student)
- PostBio2 ~ PreBio1 + PostBio1 + NonMolecular + (1|Student)
- Post2Bio2 ~ PreBio1*Molecular + PostBio1*NonMolecular + (1|Student)

**Table S5 Regression output for student responses to trait gain prompts: molecular vs. nonmolecular items**

All models included a random variable identifying individual students, to control for repeated measures. Estimates report logodds.

A. Model 1: PostBio1 ~ PreBio1Score + Topic + (1|Student)

|  | Estimate | Std.Error | z-value | *p*-value |
| --- | --- | --- | --- | --- |
| (Intercept) | -1.70 | 0.08 | -20.22 | < 0.0001 |
| PreBio1 | 0.15 | 0.0 | 6.84 | < 0.0001 |
| NonMolecular | 0.09 | 0.08 | 1.17 | 0.24 |

B. Model 2: PostBio2Score ~ PreBio1Score + Topic + (1|Student)

|  | Estimate | Std.Error | z-value | *p*-value |
| --- | --- | --- | --- | --- |
| (Intercept) | -1.71 | 0.09 | -19.71 | < 0.0001 |
| PreBio1 | 0.08 | 0.02 | 3.22 | 0.001 |
| NonMolecular | -0.33 | 0.09 | -3.57 | < 0.001 |

Controlling for scores prior to entering the course sequence, there is no difference in molecular vs. nonmolecular scores on trait gain items at the end of the initial term (Table S5A). However, controlling for their score before entering the course sequence, the logodds of scoring additional points on molecular items at the end of the series was higher than the logodds of scoring additional points on nonmolecular items at the end of the series (Table S5B, Figure S3A).

To compare understanding of molecular versus nonmolecular rubric items on trait-loss prompts, we ran the same five models as for the trait-gain analysis. The second model (PostBio2 ~ PreBio1) did not fit. We used AIC to favor model 4 (controlling for PreBio1 and PostBio1 without interaction terms) over model 5. We found that, in Bio1, students were equally likely to earn points on nonmolecular items as molecular items (Table S6A); in Bio2, students were more likely to earn points on molecular items than nonmolecular items (Table S6B). Finally, controlling for both PreBio1 and PostBio1 score, students were more likely to get a higher score on molecular than nonmolecular items on the PostBio2 survey (Table S6C).

The results of the responses to both trait-gain and trait-loss questions indicate that differences between student responses on molecular versus nonmolecular questions emerged in the second course, when the proportion of correct responses to molecular aspects of the rubric exceeded the number of correct responses to nonmolecular aspects of the rubric.

**Figure S3 Violin chart of student responses across two terms: molecular vs nonmolecular items**

A.
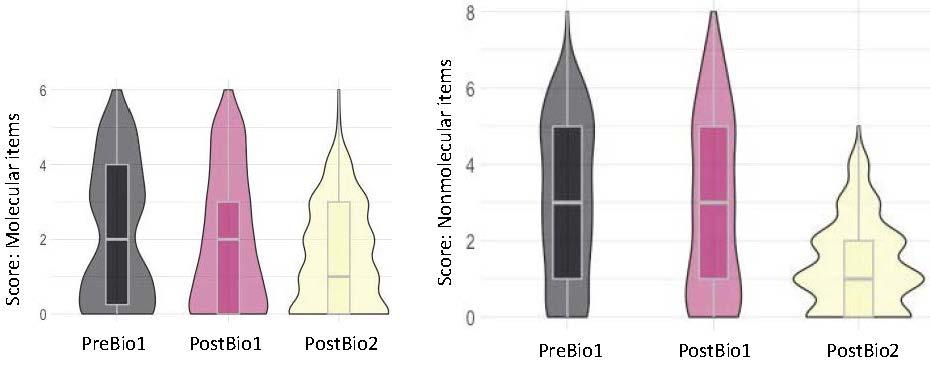
Trait gain responses
B. Trait loss responses

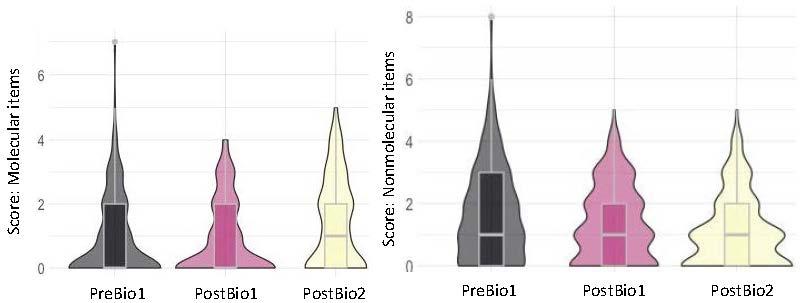

**Table S6 Regression output for student responses to trait loss prompts: molecular vs. nonmolecular items**

A. Model 1: PostBio1Score ~ PreBio1Score + NonMolecular + (1|Student)

|  | Estimate | Std.Error | z-value | *p*-value |
| --- | --- | --- | --- | --- |
| (Intercept) | -2.31 | 0.07 | -31.3 | <0.0001 |
| PreBio1 | 0.17 | 0.03 | 5.57 | <0.0001 |
| NonMolecular | 0.06 | 0.08 | 0.72 | 0.47 |

B. Model 3: PostBio2Score ~ PostBio1Score + NonMolecular + (1|Student)

|  | Estimate | Std.Error | z-value | *p*-value |
| --- | --- | --- | --- | --- |
| (Intercept) | -1.60 | 0.06 | -27.39 | <0.0001 |
| PostBio1 | 0.14 | 0.03 | 4.68 | <0.0001 |
| NonMolecular | -0.55 | 0.07 | -7.99 | <0.0001 |

C. Model 4: PostBio2Score ~ PreBio1Score + PostBio1Score + NonMolecular + (1|Student)

|  | Estimate | Std.Error | z-value | *p*-value |
| --- | --- | --- | --- | --- |
| (Intercept) | -1.97 | 0.10 | -20.32 | < 0.0001 |
| PreBio1 | 0.10 | 0.04 | 2.50 | 0.01 |
| PostBio1 | 0.16 | 0.04 | 3.57 | < 0.001 |
| NonMolecular | -0.51 | 0.11 | -4.83 | < 0.0001 |

***Research Question 4: Did performance on prompts about trait gains differ from performance on prompts about trait loss?***

Research on the ACORNS prompts showed that students have a harder time explaining the loss of traits under evolution by natural selection versus the gain of traits (Nehm and Ha, 2011). The difficulty is understandable, as in many cases the fitness advantage of losing a trait is poorly understood. Consider, for example, the loss of functional eyes in vertebrate populations that colonize caves. Biologists may explain the observation under the assumption that after a change in the environment, traits that no longer serve a positive function carry a fitness cost that favors loss-of-function mutations, but that hypothesis is seldom tested in a rigorous way.

To test the prediction that students would perform worse on trait loss prompts than trait gain prompts evaluated under the E-ACORNS rubric, we fit five models:

- Model 1: PostBio1 ~ PreBio1 + GL + (1|Student)
- Model 2: PostBio2 ~ PreBio1 + GL + (1|Student)
- Model 3: PostBio2 ~ PostBio1 + GL + (1|Student)
- Model 4: PostBio2 ~ PreBio1 + PostBio1 + GL + (1|Student)
- Model 5: PostBio2 ~ PreBio1 * GL + PostBio1 * GL + (1|Student)

where “GL” is a dummy variable indicating gain or loss responses, “(1|Student)” represents a random effect controlling for repeated measures from the same student, and “*” indicates an interaction term.

We used AIC to favor model 4 (controlling for PreBio1 and PostBio1 without interaction terms) over model 5. We find that controlling for prescore, after Bio1 the log odds of earning points on gains is higher than the logodds of earning points on loss (Table S7A). However, this pattern is not maintained in Bio2, nor when considering the entire sequence: controlling for prescore (either before Bio1 or after Bio1, or both), there was no difference in the logodds of earning points on trait-gain versus trait-loss after Bio2 (Table S7B-D). Because Loss did not have a statistically significant impact on explaining variation in PostBio2 scores controlling for earlier scores, we conclude that our prediction of loss being consistently harder was not borne out in this population. However, from inspection of the data graphed in Figure S4, we hypothesize that differences between trait-gain and trait-loss scores disappeared due to a dramatic decline in trait-gain scores during the second term. Thus, conclusions about loss being consistently harder merit further investigation.

**Table S7 Regression output for student responses on trait gain versus trait loss prompts**

A. Model 1: PostBio1Score ~ PreBio1Score + GL + (1|Student)

|  | Estimate | Std.Error | z-value | *p*-value |
| --- | --- | --- | --- | --- |
| (Intercept) | -1.74 | 0.10 | -17.83 | < 0.0001 |
| PreBio1 | 0.10 | 0.01 | 6.62 | < 0.0001 |
| Loss | -0.53 | 0.08 | -6.90 | < 0.0001 |

B. Model 2: PostBio2Score ~ PreBio1Score + GL + (1|Student)

|  | Estimate | Std.Error | z-value | *p*-value |
| --- | --- | --- | --- | --- |
| (Intercept) | -2.06 | 0.11 | -19.35 | < 0.0001 |
| PostBio1 | 0.06 | 0.02 | 3.53 | < 0.001 |
| Loss | -0.02 | 0.08 | -0.20 | 0.84 |

C. Model 3: PostBio2Score ~ PostBio1Score + GL + (1|Student)

|  | Estimate | Std.Error | z-value | *p*-value |
| --- | --- | --- | --- | --- |
| (Intercept) | -2.18 | 0.10 | -22.03 | < 0.0001 |
| PostBio1 | 0.08 | 0.01 | 5.70 | < 0.0001 |
| Loss | 0.10 | 0.08 | 1.14 | 0.25 |

D. Model 4: PostBio2Score ~ PreBio1Score + PostBio1Score + GL + (1|Student)

|  | Estimate | Std.Error | z-value | *p*-value |
| --- | --- | --- | --- | --- |
| (Intercept) | -2.30 | 0.12 | -19.80 | < 0.0001 |
| PreBio1 | 0.03 | 0.02 | 1.88 | 0.06 |
| Post Bio1 | 0.07 | 0.021 | 4.74 | < 0.0001 |
| GL Loss | 0.15 | 0.09 | 1.70 | 0.09 |

**Supplemental Literature cited**

Anderson D, Burnham K (2004). Model Selection and Multi-Model Inference. NY: Springer-Verlag.

Koo TK, Li MY (2017) A guideline of selecting and reporting intraclass correlation coefficients for reliability research. Journal of Chiropractic Medicine 16: 346-355.

R Core Team (2021). R: A language and environment for statistical computing. R Foundation for

Statistical Computing, Vienna, Austria; [https://www.R-project.org/](https://www.r-project.org/).

Theobald E (2018) Students are rarely independent: when, why, and how to use random effects in discipline-based education research. CBE-Life Sciences Education 17: rm2, 1-12.

Theobald EJ, Aikens M, Eddy S, Jordt H (2019) Beyond linear regression: A reference for analyzing common data types in discipline-based education research. Physical Review Physics Education Research 15, 020110.
